# Deep Learning model accurately classifies metastatic tumors from primary tumors based on mutational signatures

**DOI:** 10.1101/2022.09.29.510207

**Authors:** Weisheng Zheng, Mengchen Pu, Xiaorong Li, Sutong Jin, Xingshuai Li, Jielong Zhou, Yingsheng Zhang

## Abstract

Metastatic propagation is the leading cause of death for most cancers. Prediction and elucidation of metastatic process is crucial for the therapeutic treatment of cancers. Even though somatic mutations have been directly linked to tumorigenesis and metastasis, it is less explored whether the metastatic events can be identified through genomic mutation signatures, a concise representation of the mutational processes. Here, applying mutation signatures as input features calculated from Whole-Exome Sequencing (WES) data of TCGA and other metastatic cohorts, we developed MetaWise, a Deep Neural Network (DNN) model. This model accurately classified metastatic tumors from primary tumors. Signatures of non-coding mutations also have a major impact on the model performance. SHapley Additive exPlanations (SHAP) and Local Surrogate (LIME) analysis into the MetaWise model identified several mutational signatures directly correlated to metastatic spread in cancers, including APOBEC-mutagenesis, UV-induced signatures and DNA damage response deficiency signatures.

## Introduction

For most cancers, the metastatic spread is the main cause of cancer morbidity and mortality^1^. It’s been reported that metastasis is responsible for 90% of cancer deaths^2^. Once detaching from the primary site, metastatic cancer cells can invade all parts of the body through the circulating system^3,4^. However, the distribution of metastatic sites for a given primary tumor is not random and is dictated by factors such as the anatomical location, the cell of origin, and molecular subtypes, among others^5^. The identification of metastatic tumors and their tumor of origins is pivotal to therapeutic treatment in clinical practice.

Somatic mutations accumulate throughout the lifetime of an individual^6,7^. The majority of these somatic mutations are neutral and increase in a passive manner, while some genomic changes on the DNA sequences do affect the regulation and function of genes, leading to the abnormal phenotypes of cells^6^. Accumulation of mutations in key regulatory genes can lead to the genesis of diseases, such as cancers. Therefore, mutations as a feature set can be an informative candidate to decipher the developmental stages of tumorigenesis, such as, primary and metastatic tumors.

Aberrant somatic mutations in DNA take many different forms, including single base substitutions (SBS), doublet base substitutions (DBS), small insertions and deletions (ID), and others. These genomic changes result from diverse processes including replication infidelity, spontaneous or enzymatic conversions and exposure to exogenous or endogenous mutagens^8^. Each of these processes leaves the tumor genome with a distinctive pattern of mutations known as mutational signatures^9^. Mathematical extraction of mutational signatures in large pan-cancer series revealed more than 100 different signatures, including SBS^10–12^, DBS^12,13^ and ID signatures^13^. While the mechanism of many of the signatures are still unknown, some of them are linked to established (such as smoking or UV radiation exposure) or novel (such as APOBEC mutagenesis) etiologies. Whole-Genome Sequencing (WGS) analysis of the metastatic solid tumors in pan-cancers has revealed the specific patterns of mutational signatures in metastatic tumors^14^. Analysis of the matched primary and metastatic tumors further demonstrated the transformed profiles of mutational signatures from primary to metastatic tumors in colorectal^15^ and papillary thyroid cancers^16^. Additionally, based on SBS mutational signature, Sina Abdollahi, et al. distinguished later stage cancers from tumors of other stages using a feed-forward neural network^17^.

Here, we propose that SBS along with DBS and ID mutational signatures as a characteristic feature in cancer genome can be used as informative probes to distinguish primary and metastatic cancers. To test this hypothesis, we performed mutational signature analysis of primary and metastatic tumors from TCGA pan-cancer studies containing more than 9000 primary and several metastatic tumor cohorts with more than 1500 metastatic tumors. By including these features to the MetaWise model, we can accurately classify primary and metastatic tumors in the training, validation and test sets (Fig. 1). Comparing with existing model^17^, our model has a better performance in the identification of metastatic tumors from primary tumors with an averaged accuracy of 86% on the test sets. Furthermore, we extracted several signatures that are the most informative for our model using SHAP^18^ and LIME^19^ methods (Fig. 1). These signatures, responsible for the growth of cancerous metastatic cells, include signatures linked to APOBEC and UV exposure, among others.

**Figure 1.**
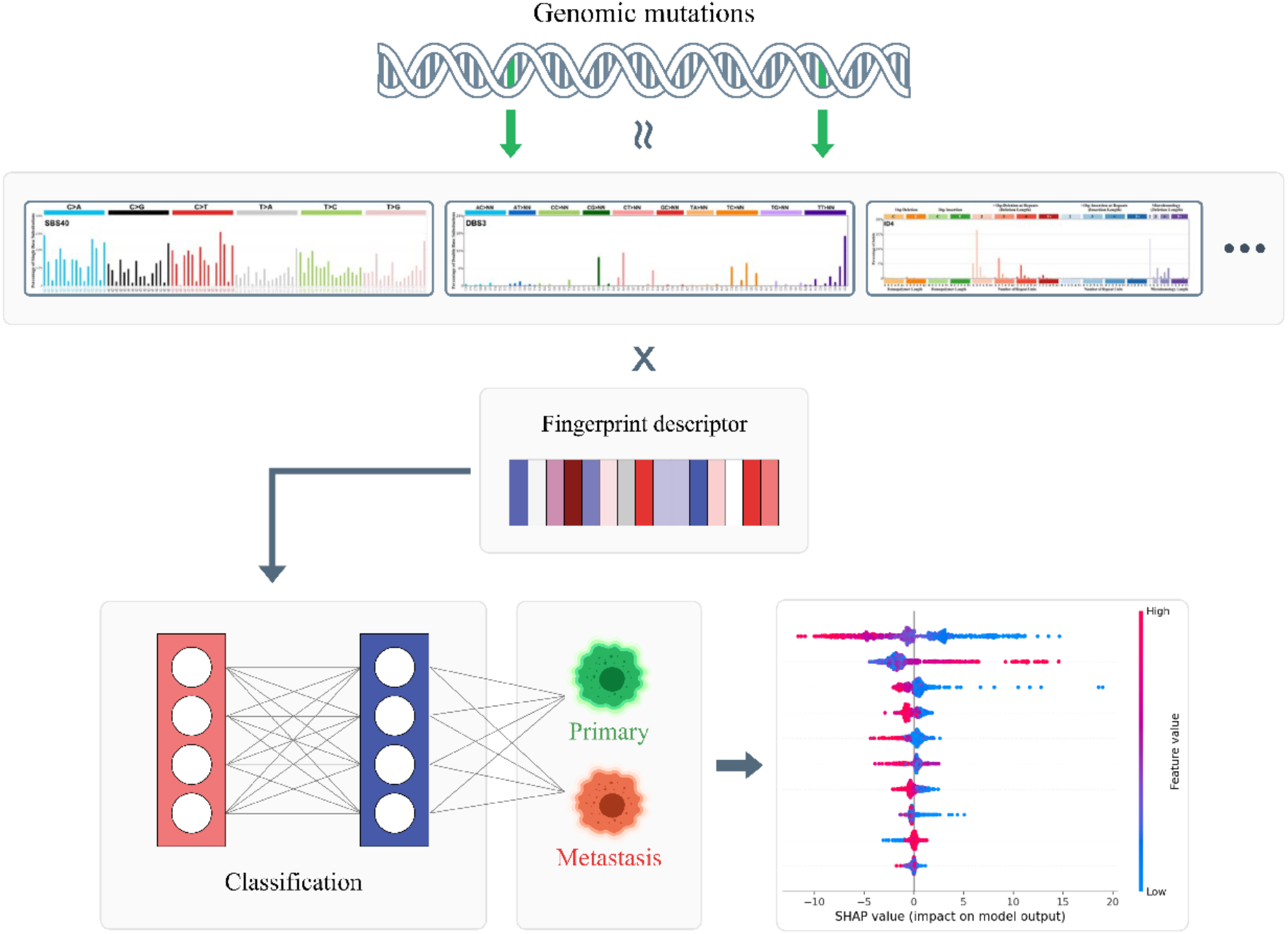
MetaWise model workflow. Somatic mutations detected by WES or WGS data are characterized as SBS, DBS and ID mutational signatures and the calculated signatures in each sample are used as the input feature to the MetaWise model. MetaWise classify the sample as primary or metastatic tumors based on the mutational signatures fingerprint descriptor. Moreover, SHAP and LIME are applied to extract the most informative signatures for the model.

## Results

### Mutational landscape of primary and metastatic tumors

The Cancer Genome Atlas (TCGA) is a landmark pan-cancer project containing somatic mutations, copy number variations, gene expression profiles and other genomic characteristics among over 10000 tumors from more than 30 different cancer types^20^. In order to study the mutational landscapes of primary tumors and metastatic tumors, and construct data sets for Deep Learning (DL) models, we retrieved the somatic mutations detected from WES of TCGA project^20^ and several metastatic tumor cohorts, such as MET-500^21^, BRCA-igr^22^, SKCM-dfci^23^, and others^24–26^ from cBioPortal (https://www.cbioportal.org/) database^27^. These data sets consist of more than 9700 primary tumors from more than 25 different cancer types and about 1500 metastatic specimens from more than 30 cancer types (Fig. 2a, b). The broad spectrum of cancer types in these data sets allows us to train a more general and un-biased DL model for tumor stages classification.

**Figure 2.**
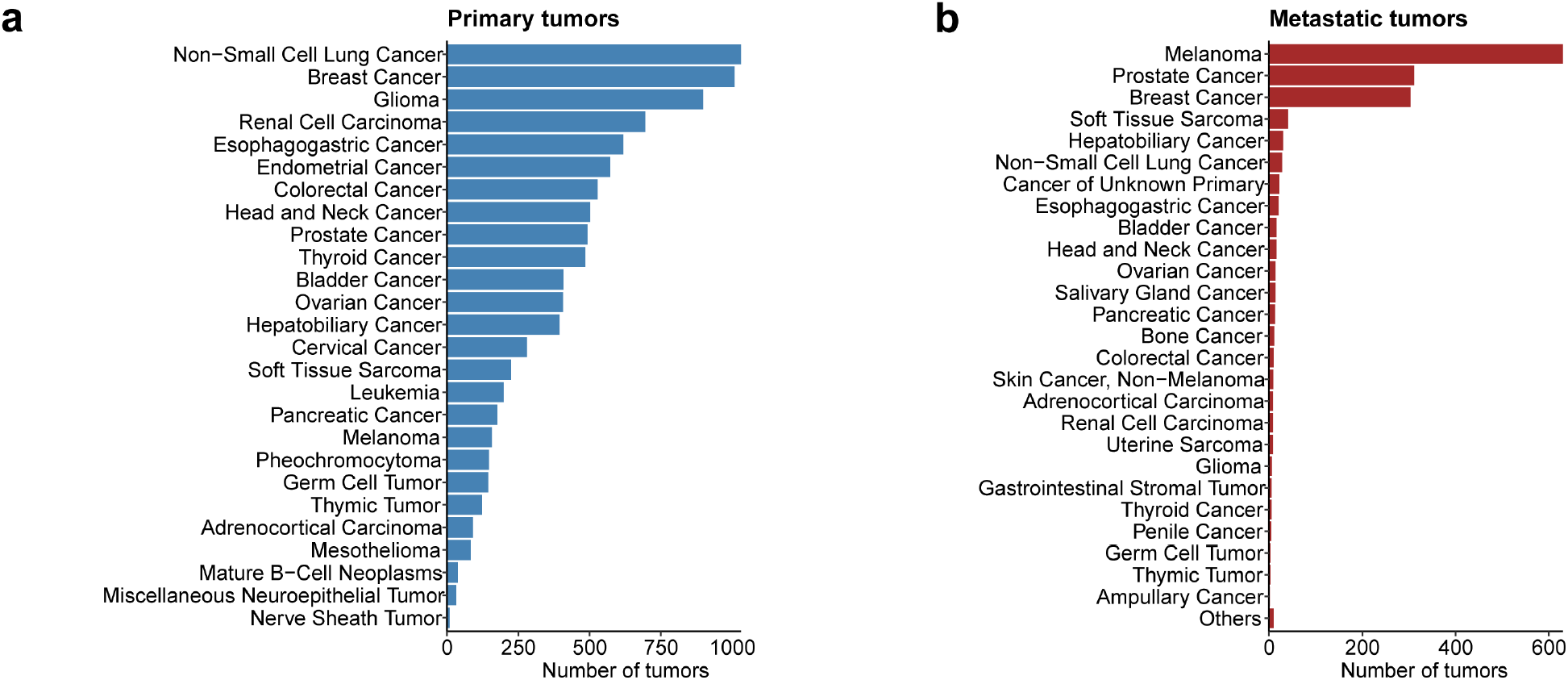
Cancer type distribution of our datasets. **a.** The number of tumors in different cancer types of primary tumors from TCGA pan-cancer cohorts. **b.** The number of metastatic tumors in different cancer types from TCGA and several other metastatic tumor cohorts.

To study the mutational landscape, we conducted mutational signature analysis of the somatic mutations from primary and metastatic tumors (Methods). Using the same procedures described in previous study^17^, we extracted 30 *de novo* SBS mutational signatures from these data using SigProfiler^13,28^. The Catalogue Of Somatic Mutations In Cancer (COSMIC) website has also curated 78 SBS, 11 DBS and 18 ID mutational signatures, some of which associated with known etiologies (V3.3)^13^. We believe that inclusion of DBS and ID signatures to the data sets can provide extra information, and adopted COSMIC mutational signatures as references to calculate the exposure matrix offer reliable information on the performance of the model. Applying difference of relative frequency (DRF) analysis^17^ to the 30 *de novo* SBS mutational signatures and 107 COSMIC signatures profile, we revealed significant difference for some signatures between primary and metastatic tumors (Fig. 3a, b). The DRF analysis indicates that UV-induced signatures SBS7a and SBS7b as well as chemical treatment induced signatures SBS11 are enriched in metastatic tumors (Fig. 3b).

**Figure 3.**
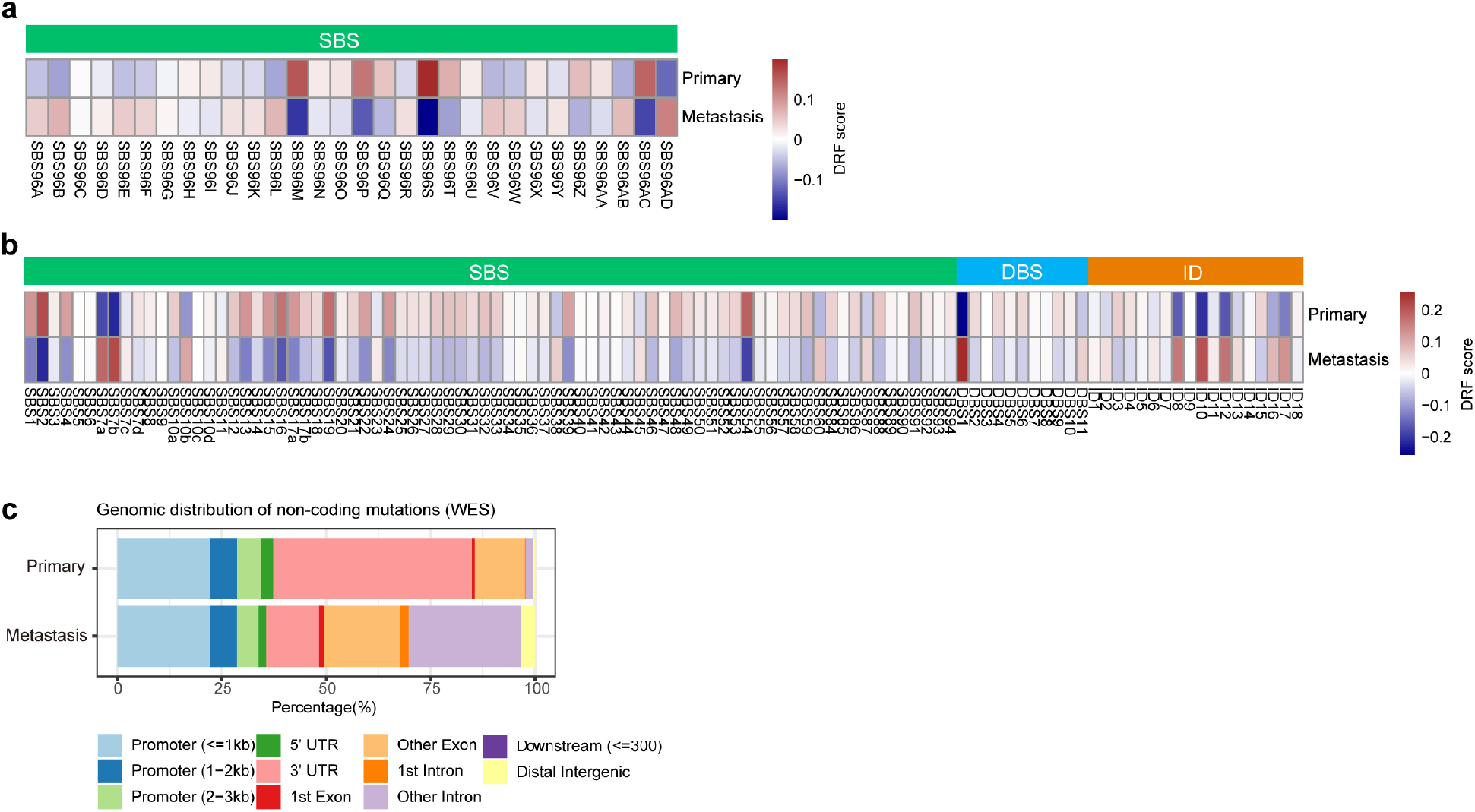
Mutation landscapes in primary and metastatic tumors. **a.** The DRF results calculated from 30 *de novo* SBS signatures. **b.** The DRF results calculated from 107 COSMIC mutational signatures. **c.** The genomic distribution of non-coding mutations detected by WES data in primary and metastatic tumors.

### MetaWise model accurately classify metastatic tumors from primary tumors based on mutational signatures

The different distributions of mutational signatures between primary and metastatic tumors lay the foundation that mutational signatures can be used as input features in DNN models to classify metastatic tumors from primary tumors (Fig. 3a, b). In order to verify this, we trained the MetaWise, a ten-layered fully connected DNN model, employing mutational signatures profiles to classify metastatic and primary tumors using 5-fold cross validation on each data set. Due to the imbalanced population of primary and metastatic tumors, we randomly split the primary tumors into 5 groups to match the number of metastatic tumors. First, we trained the DNN model based on the 30 *de novo* SBS signatures profile alone. The averaged accuracies of the predictions are 78.65% and 78.37% on validation and test set, respectively (Table 1). These outcomes are comparable with the prediction results of DiaDeL model from the earlier study which use the same SBS signature profiles as input^17^.

**Table 1.** The model performance on different dataset.

|  | Dataset | Training accuracy | Validation accuracy | Test accuracy |
| --- | --- | --- | --- | --- |
| MetaWise | WGS data | 98% | 96.8% | 94.2% |
| MetaWise | WES cBioportal 30 | 91.36% | 78.65% | 78.37% |
| MetaWise | WES cBioportal 107 | 92.58% | 89.16% | 88.81% |

The characterization of SBS mutational signatures alone is insufficient to represent the entire mutational landscape. Some of DBS and ID signatures have shown significantly different distributions between primary and metastatic tumors (Fig. 3b). Therefore, DBS and ID signatures are tested to improve the model performance. We trained MetaWise model based on the 107 COSMIC signature profile covering all the SBS, DBS and ID signatures. The averaged accuracies improved to 89.16% and 88.81% on validation and test set, respectively, which are significant comparing to using only 30 *de novo* SBS signatures (Table 1). These results demonstrate that DBS and ID signatures have big impacts on the mutational profiles and resulted in the improvement of the model performance. Thus, MetaWise based on the mutational signature profile can accurately classify primary and metastatic tumors.

### Non-coding mutations impact model performance

Non-coding mutations have gained significant attention recently for their contribution in tumorigenesis^29,30^. In a recent work, Wei Jiao, et al. have developed a DNN model to accurately classify cancer types based on somatic passenger mutations, most of which are non-coding mutations^31^. It is interesting to know the scope of non-coding mutations’ effects on the mutational signature profile and model performance on primary and metastatic tumor classification. There’re some non-coding mutations in the WES data sets, including variants located in the untranslated regions (UTRs), introns, and other regions (Fig. 3c). To test the influence of these mutations, we subtracted the non-coding mutations from the data sets, and conducted mutational signature analysis on these data as described above. We tested MetaWise based on the mutational signature profiles with and without non-coding mutations. The results show that MetaWise with either 30 *de novo* SBS signatures or 107 COSMIC signatures profiles consistently has higher performance based on the data sets with non-coding mutations (Table 2), which demonstrate the significance of non-coding mutations in the characterization of mutational signatures and DNN model performance.

**Table 2.**
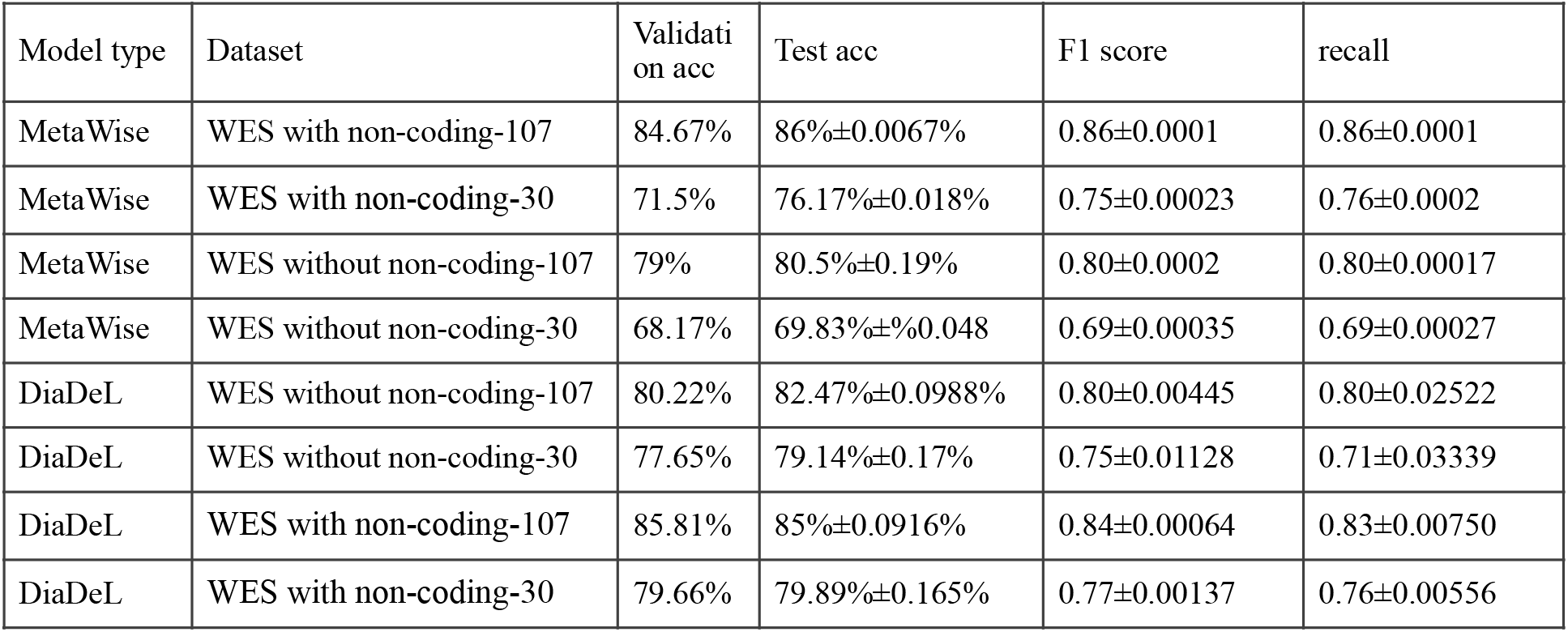
The performance comparison between MetaWise and DiaDeL model on data sets with and without non-coding mutations.

| Model type | Dataset | Validation acc | Test acc | F1 score | recall |
| --- | --- | --- | --- | --- | --- |
| MetaWise | WES with non-coding-107 | 84.67% | 86%±0.0067% | 0.86±0.0001 | 0.86±0.0001 |
| MetaWise | WES with non-coding-30 | 71.5% | 76.17%±0.018% | 0.75±0.00023 | 0.76±0.0002 |
| MetaWise | WES without non-coding-107 | 79% | 80.5%±0.19% | 0.80±0.0002 | 0.80±0.00017 |
| MetaWise | WES without non-coding-30 | 68.17% | 69.83%±0.048% | 0.69±0.00035 | 0.69±0.00027 |
| DiaDeL | WES without non-coding-107 | 80.22% | 82.47%±0.0988% | 0.80±0.00445 | 0.80±0.02522 |
| DiaDeL | WES without non-coding-30 | 77.65% | 79.14%±0.17% | 0.75±0.01128 | 0.71±0.03339 |
| DiaDeL | WES with non-coding-107 | 85.81% | 85%±0.0916% | 0.84±0.00064 | 0.83±0.00750 |
| DiaDeL | WES with non-coding-30 | 79.66% | 79.89%±0.165% | 0.77±0.00137 | 0.76±0.00556 |

To directly compare our model with state-of-the-art model DiaDeL, we built the model based on the same datasets above. Both DiaDeL and MetaWise perform significantly better using the mutational signatures containing non-coding mutations than those without non-coding mutations (Table 2). Besides, MetaWise in the COSMIC signatures with non-coding mutations has the highest performance (Table 2, with about 86% testing accuracy, 0.86 F1 score and 0.86 recall), showing the great power of this DNN model with the inclusion of SBS, DBS and ID mutational signatures from both coding and non-coding regions.

WGS detects genomic alterations on a whole genome scale, thus most of the alterations in the non-coding regions can be revealed. To further investigate the effect of non-coding mutation signatures beyond the UTR and intron regions on model performance, we studied the somatic mutations detected by WGS from the Pan-Cancer Analysis of Whole-Genome (PCAWG) project^32^, including more than 1600 primary and 100 metastatic tumors. We randomly selected roughly 100 primary tumors and 100 metastatic tumors from the WGS data of PCAWG and constructed the WGS dataset for our model testing. Applying MetaWise model to this ‘few-shot’ WGS data, we get a classification result with very high accuracy (averaged accuracy 96.8% and 94.2% on validation and test sets) (Table 1). Taken together, these results demonstrate that noncoding mutational signatures from whole genome greatly improve the mutational landscape of cancer genome and the model performance.

### Feature selection to interpret the MetaWise model

DRF analysis of mutational signatures distribution revealed the significant difference of some mutational signatures between primary and metastatic tumors (Fig 3 a, b). These differences are likely to give the distinguishing power to the MetaWise model. To decipher which mutation signatures contribute to the performance of the model, we executed two feature selection methods, SHAP and LIME, respectively. The SHAP analysis performs hyper-parameter tuning and feature selection simultaneously in a single pipeline^18^, while LIME analysis performs feature selection via training local surrogate models^19^. The results of SHAP show that clock-like signature SBS1, APOBEC associated signature SBS2, UV-light induced signatures (SBS7a/ SBS7b/DBS1) and several ID signatures are the most informative mutational signatures for our model (Fig. 4). LIME analysis further showed that DNA repair associated signatures (SBS3/ SBS36/SBS44/ID8) and APOBEC associated signature SBS2 are the most informative signatures for the classification of primary and metastatic tumors (Fig. 4). It’s been reported that APOBEC mutagenesis is associated with tumor evolution and heterogeneity in metastatic thoracic tumors^33^. Besides, UV-radiation induced inflammation promotes the metastasis in melanoma^34^. In addition, DNA damage response deficiencies, such as defective of homologous recombination DNA damage repair (SBS3) and defective of DNA mismatch repair (SBS44), have been reported to be enhanced in the brain metastasis of colorectal cancer^15^. Taken together, these results demonstrate these mutational signatures can have great contribution to the metastatic event in cancers.

**Figure 4.**
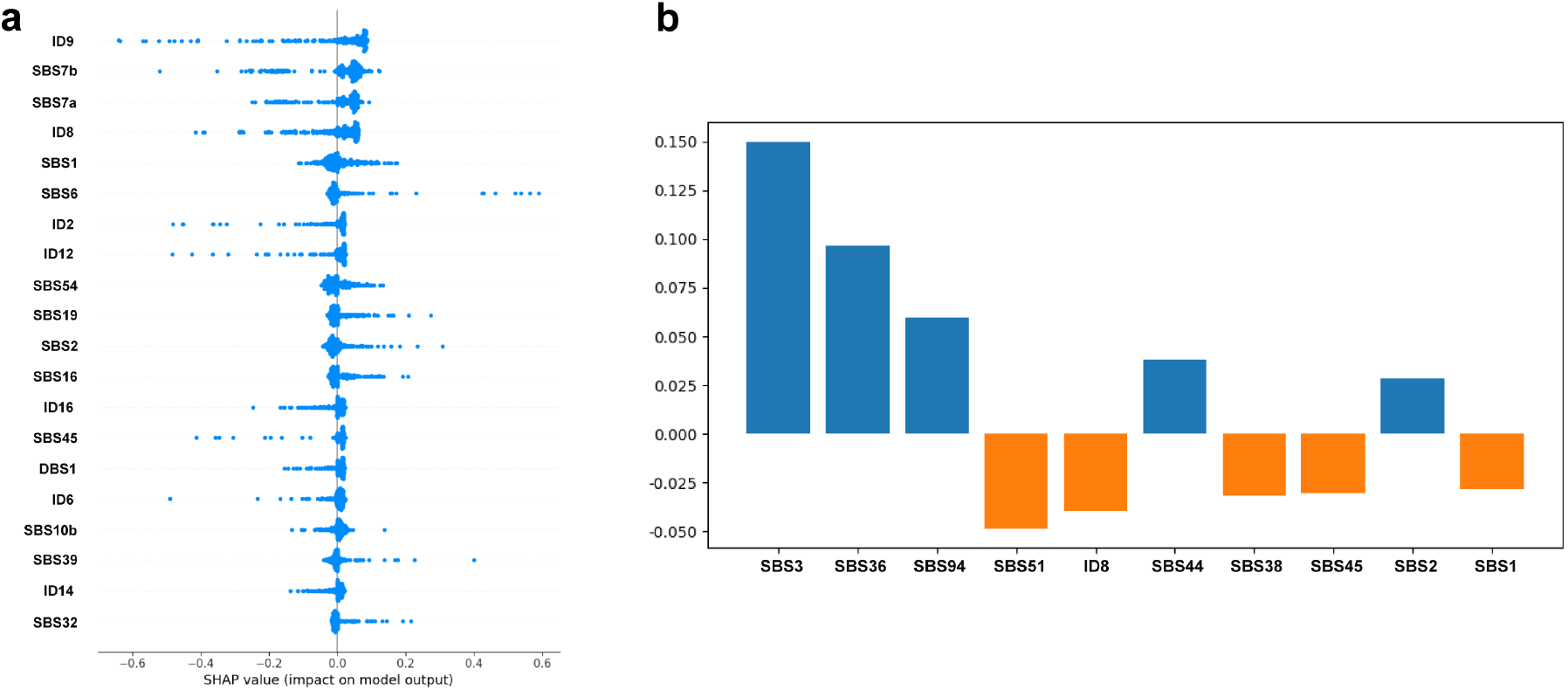
Interpreting the MetaWise model with the framework of SHAP and LIME. The results show the most informative mutational signatures selected by SHAP (**a.**) and LIME (**b.**).

## Discussion

To design DNN models to distinguish metastatic tumors from primary tumors, we show that combining SBS with DBS and ID signatures as input features significantly improved the model performance. Compared to SBS signatures, DBS and ID signatures are less explored due to the low frequency in cancer genome and the limitations in characterization methods. With the development of high-throughput sequencing on cancer genome, numerous DBS and ID signatures have been identified^12,13^. The genomic distribution and sequence compositions of DBS and small IDs are non-random^10,11^ and also associated with known mutational processes^12,13^. Therefore, DBS and ID signatures can provide insightful biological relevant information to the models. Recently, Ruben M.Drews, et al.^35^ and Christopher D Steele, et al.^36^ have introduced two types of characterization of chromosomal instability (CIN) signatures through pan-cancer studies. It’s been reported that chromosomal instability results in metastasis through the cGAS-STING cytosolic DNA-sensing pathway^37^. In future work, it is desirable to include CIN signatures in metastatic tumor predicting models.

In our results, inclusion of non-coding mutations in the mutation profiles of WES data improve the characterization of mutational signatures and model performance. Most of the noncoding mutations detected by WES data locate in the key regulatory regions, such as promoters and UTRs. Interestingly, there’re much more proportion of mutations occurred in the 3’ UTR regions in primary tumors than metastatic ones (Fig. 3c). Regulatory elements, such as miRNA, regulate gene function through targeting the 3’ UTR regions of the mRNA transcripts^38^. The somatic mutations in these regions might affect the post-transcriptional regulation of key regulatory genes, leading to tumorigenesis and metastatic spread of cancers^39,40^. Although we were unable to test WGS data on a larger scale due to the data availability. Using a few-shot approach of WGS data, the performance of the model demonstrated a high degree of improvement, indicating the importance of the non-coding mutations profiles and the necessary to include a larger cohort of WGS data.

The DRF analysis identified significant difference in mutational signatures distribution between primary and metastatic tumors, including UV-induced signatures and others. Applying SHAP and LIME analysis to the MetaWise model, we obtained informative mutational signature combination to distinguish metastatic tumors from primary tumors, which are in concordance with the results of DRF analysis. Signatures of APOBEC-, UV-induced and DNA damage response deficiency are the most significant mutational processes in tumor metastasis^33,34^, which deserve further investigation in primary to metastatic tumor transition processes.

## Methods

### Data sets

The somatic mutations of WES dataset of TCGA and other metastatic cohorts were downloaded from cBioPortal database (https://www.cbioportal.org/)^27^ with cBioPortalData packages in R environment^41^. The WES data were constructed into two data sets, one with the whole somatic variants in the coding and non-coding regions, and the other with only the variants detected in the coding regions. These two data sets were used to study the influence of non-coding variants in the model performance. We also obtained somatic mutations of WGS data from the PCAWG project (https://dcc.icgc.org/pcawg)^32^. The WGS data is consist of more than 1600 primary tumors and about 100 metastatic tumors.

### Mutational signature analysis

The single-base substitutions were classified into 96 SBS categories considering the 6 substitution types (C>A, C>G, C>T, T>A, T>C and T>G) and their 5’ and 3’ adjacent bases. The doublet bases substitutions were classified into totally 78 DBS categories considering the doublet bases substitution types (AC>NN, AT>NN, CC>NN, CG>NN, CT>NN, GC>NN, TA>NN, TC>NN, TG>NN and TT>NN) and their flanking bases. The small insertion and deletion mutations were classified into 83 ID categories as previous reported considering the insertion or deletion types and the number of repeat lengths^13,42^. In order to test DiaDeL model, we extracted 30 *de novo* SBS mutational signatures with 96 SBS categories using SigProfiler^13,28^ (Supplementary Table 1). We downloaded SBS, DBS and ID signatures from COSMIC v3.3 and used Sigminer^43^ to fit the activities of these signatures in the primary and metastatic tumors. The exposure matrices of 30 *de novo* SBS mutational signatures and the 107 COSMIC mutational signatures were used as input features for DL models (Supplementary Table 2, 3).

### Difference of relative frequency (DRF) analysis of mutational signatures

The DRF analysis of mutational signatures is adjusted from previous study^17^. We used an equation as below:

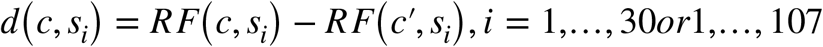

Where *s_i_* represents *i^th^* signature and *RF*(*x, s_i_*) represents the relative frequency of signature *s_i_* in primary or metastatic tumors. If *RF*(*c, s_i_*) represents the relative frequency of signature *s_i_* in primary tumors then *RF*(*c*′, *s_i_*) represents the relative frequency of *s_i_* in the metastatic tumors, and vice versa.

### Neural network architecture and training procedure

In this research, we proposed a ten-layered fully connected DNN model to differentiate the mutation profiles between primary and metastatic tumors based on multiple cancer datasets. Furthermore, we applied an existing method (DiaDeL model)^17^ for comparison.

The model is implemented in Keras using Tensorflow^44^. Mutational signature selection, parameter tuning and model training were performed. The learning rate, weight decay, dropout layer and activation method were fine-tuned using Bayesian optimization^45^.

Due to the imbalanced population of datasets, we randomly divided the primary tumors in to five groups. The performance metrics calculated in this experiment took into consideration the predictions for the validation or test sets after a rigorous internal 5-fold cross-validation. For comparability of results, we used the same split of data for both models. In addition, We apply the interpretative machine learning techniques, SHAP^18^ and LIME^19^, to decipher the significance of different mutation signatures on the prediction.

## Supporting information

Supplemental Table1_SigProfiler_30-denovo-sigs

## Data Availability

All somatic mutations detected by WES of primary and metastatic tumors from TCGA project and other metastatic cohorts are available in cBioPortal database (https://www.cbioportal.org/). The somatic mutations detected by WGS data of PCAWG are available in PCAWG website (https://dcc.icgc.org/pcawg). The processed data in our study can be retrieved from supplementary tables.

## Code Availability

The code developed for mutational signatures analysis and CNN model training and testing are available from GitHub: https://github.com/promethiume/MetaWise

## Acknowledgements

We thank G.Peng, B.Fu, Y.Xin, L.Wei and others from the Innovation Center of StoneWise, AI. Inc. for their helpful discussions and support; X.Xiang, S.Guo, Y.Wang and our colleagues at StoneWise, AI. Inc. for their support and encouragement. We thank the TCGA consortium and their foundational data by the TCGA Research Network: https://www.cancer.gov/tcga.

## Ethics declarations

### Competing interests

The authors declare no competing interests.

## Supplementary Information

Supplementary Information

## Notes

### Competing Interest Statement

The authors have declared no competing interest.

